# Extensive translation of small ORFs revealed by polysomal ribo-Seq

**DOI:** 10.1101/002998

**Authors:** Julie L. Aspden, Ying Chen Eyre-Walker, Rose J. Philips, Michele Brocard, Unum Amin, Juan Couso

## Abstract

Thousands of small Open Reading Frames (smORFs) encoding small peptides of fewer than 100 amino acids exist in our genomes. Examples of functional smORFs have been characterised in a few species but the actual number of translated smORFs, and their molecular, functional and evolutionary features are not known. Here we present a genome-wide assessment of smORF translation by ribosomal profiling of polysomal fractions. This ‘polysomal ribo-Seq’ suggests that smORFs are translated at the same level and in the same relative numbers (80%) as normal proteins. The smORF peptides appear widely conserved, show activity in cells, and display a putative amino acid signature. These findings reinforce the idea that smORFs are an abundant and fundamental genome component, displaying features usually attributed to canonical proteins, including high translation levels, biological function, amino acid sequence specificity and cross-species conservation.

---

Small Open Reading Frames (smORFs) of fewer than 100 amino acids exist in genomes in the hundreds of thousands. Bioinformatic approaches predict that hundreds if not thousands of these are translated and functional in bacteria (*1*), yeast (*2*, *3*), plants (*4*) and in animals such as *Drosophila* (*5*), mouse (*6*) and humans (*7*). This is in contrast to their low rate of detection in biochemical (proteomic) and functional (genetic) screens. A handful of cases of translated smORFs encoding bioactive peptides have been described (*8*-*11*). However, translation and functionality of smORFs at a genomic scale has not been assessed in any species and therefore the actual number of translated smORFs is not known.

**FIGURE 1.**
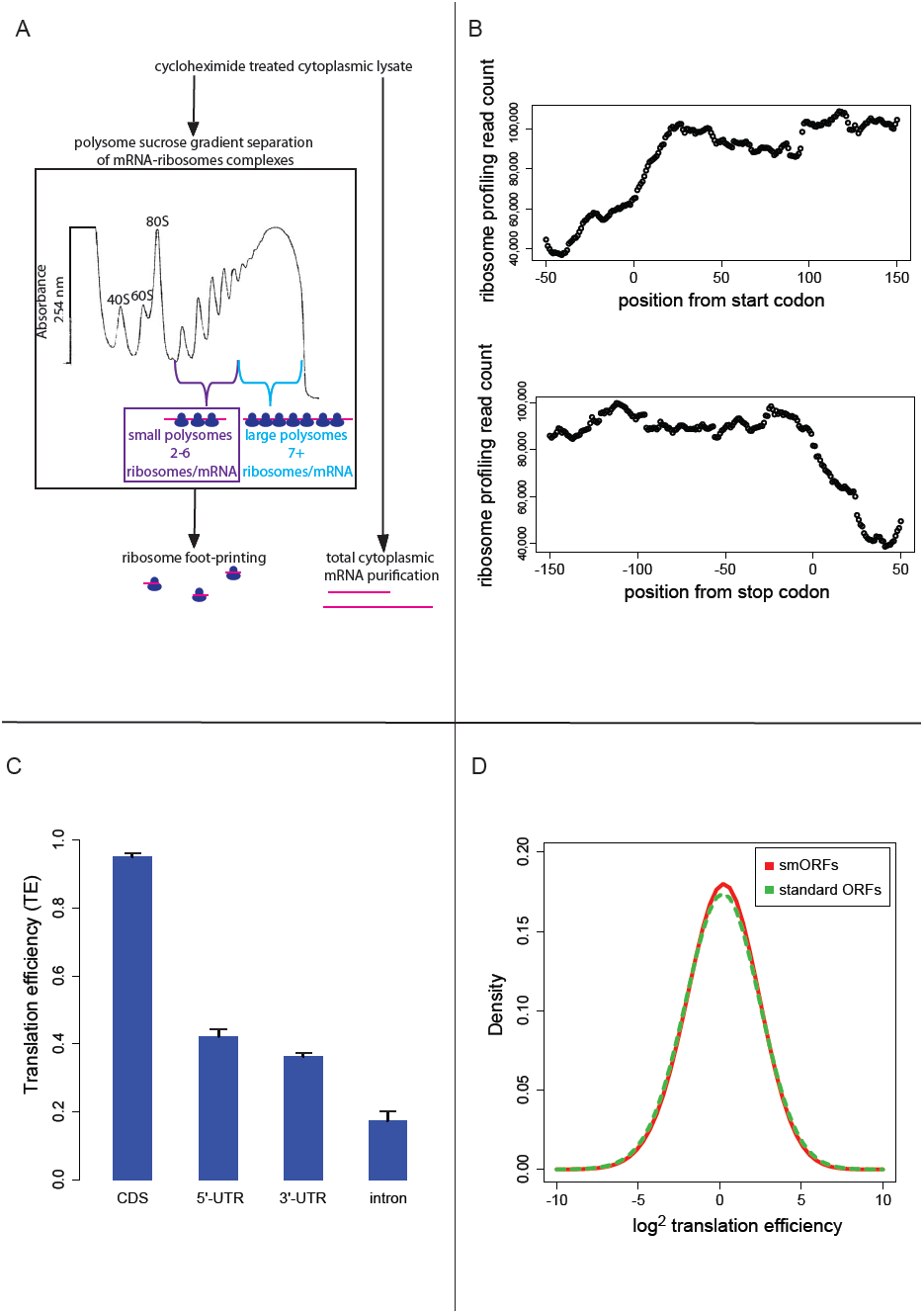
Polysomal ribo-Seq of small and large polysomes. (A) Schematic of deep polysome profiling with representative UV absorbance profile for sucrose density gradient. (B) Composite plot from all FlyBase protein-coding genes, of deep polysome profiling read counts across mRNAs in the vicinity of start (upper) and stop codons (lower) in small polysomes. (C) Median translation efficiencies of CDS, UTR and intron regions for all protein-coding genes, error bars represent SD. (D) Density plot of log^2^ translation efficiency (TE) comparing smORFs (red) and standard ORFs (green).

The new technique of ribosomal profiling has corroborated and expanded the proteomes of yeast (*12*) mouse (*13*) and zebra fish (*14*). Thousands of new translated sequences have been described in each case, whether novel exons of annotated genes, alternative initiation sites, or entire ORFs. However, the application of ribosomal profiling outside canonical translated sequences can lead to differing conclusions (*14*, *15*). The problem remains that a ribosomal footprint cannot be equated with translation; non-productive binding of single ribosomes to mRNAs and 40S ribosomal subunit scanning can result in footprints yet do not constitute translation. It has been suggested that some smORFs associate with ribosomes in such a non-productive manner and do not undergo productive translation (*16*). Moreover, smORF mRNAs are short and present a small target for ribosomal binding and generation of footprints. To overcome these difficulties, we devised an improvement to ribosomal profiling. Instead of profiling all ribosomal-bound mRNAs, small and large polysome fractions were separated, RNase-digested, and ribosome-protected mRNA fragments were deep sequenced (Fig. 1A; methods). In this way, mRNAs bound by multiple ribosomes and hence actively translated can be isolated and distinguished from mRNAs bound by sporadic, putatively non-productive single ribosomes or ribosomal subunits. This novel combination of polysome fractionation and ribosome profiling we term ‘deep polysome profiling’.

Annotated smORFs located in transcripts devoid of a canonical long coding sequence provide the most straightforward test for smORF functionality. We have assessed them in *Drosophila* S2 cells (*17*), because the *Drosophila* genome includes a high proportion of smORF genes (some 829 smORF genes, or 5% of the total) and this cell line provides enough reproducible material. To enrich for actively translating smORF mRNAs we took advantage of the limited space within a smORF (≤303 nt). Ribosomes can reach densities of 1 ribosome every 80 nt (*18*), therefore on a smORF the maximum number of ribosomes associated would be 6, allowing for 5 ribosomes in the ORF, and 1 in the 5’-UTR, scanning. RT-PCR confirmed that smORF mRNAs were enriched in small polysomes compared to large polysomes (Fig. S1A). However, small polysomes (2-6 ribosomes/mRNA) also contain mRNAs for longer ORFs being translated at less than maximum level.

Deep polysome profiling captured regions of active translation with ∼80% of reads mapped to coding sequences (Fig. 1B), their frequency dropping off before the start and after the stop codons. The translation level of individual coding sequences was provided by ribosomal density (RPKM) (*12*) and likelihood that the whole ORF is translated, by coverage of the ORF by footprints. For genes we called translated, we required ribosome density to be above 7.7 and footprint coverage to be above 0.57, which are both above the 90 percentile of the values obtained for 3’-UTRs (Table S1). These cut-offs are more stringent than previous ribosomal profiling experiments and standard RNAseq practice, and their combination should provide a robust identification of genes that undergo active translation. To overcome the possible dependence of ribosome density on RNAseq efficiency (*15*), we also used the relative metric translational efficiency (TE, RPKM of ribosome footprints/RPKM of total mRNA control reads) (*12*). We observed that median TE was significantly higher in ORFs compared to UTRs and introns (Fig 1C). 5’- and 3’-UTRs showed comparable levels of TE after we removed putative upstream open reading frames (uORFs) from 5’-UTRs. As reported for ribosomal profiling, we observe triplet phasing in the mapping of our deep polysomal reads, reflecting the positioning of ribosomes on codons (Fig. S1B).

**FIGURE 2.**
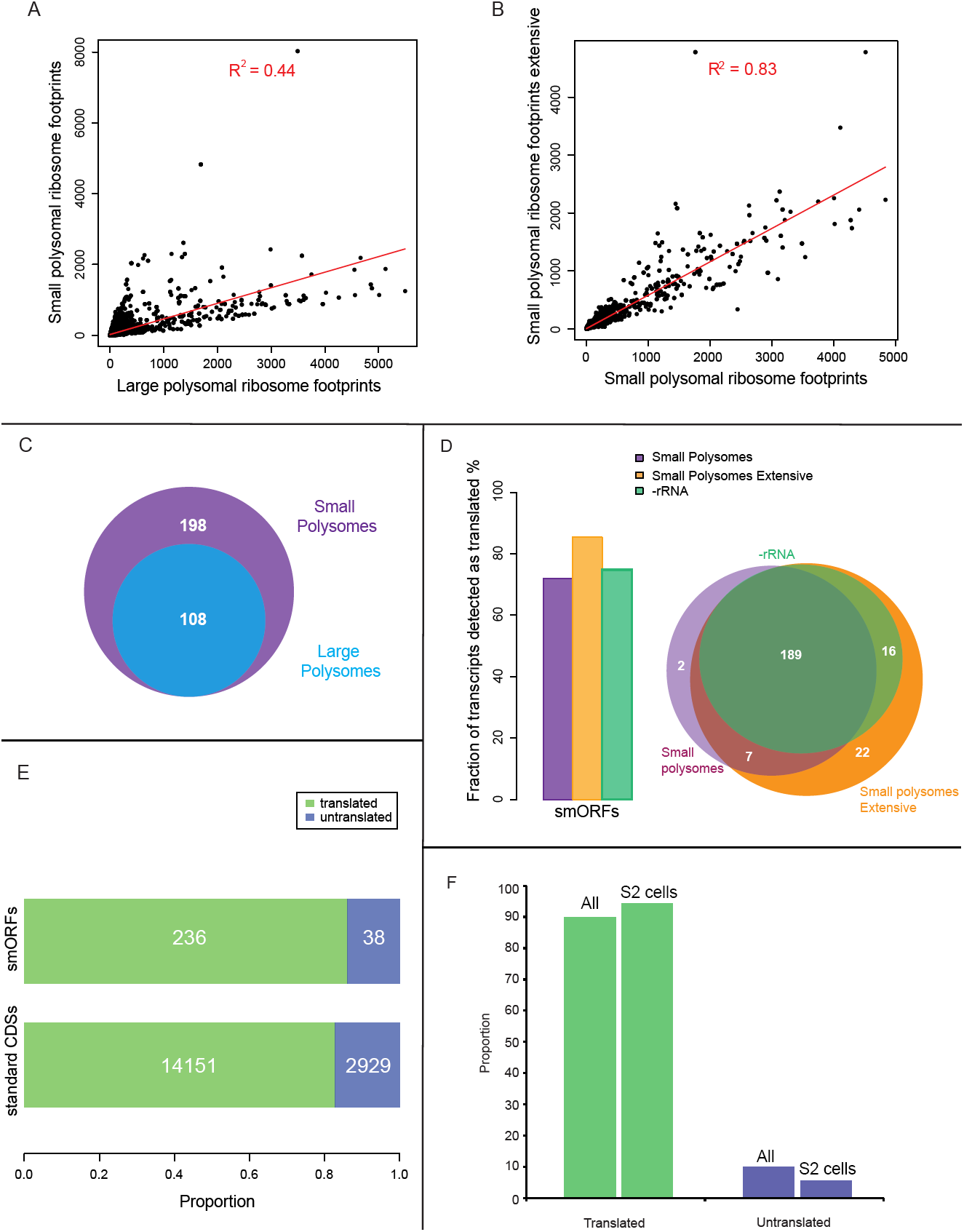
Polysomal ribo-Seq reveals translation of 236 smORFs. (A) Ribosome footprinting densities (RPKM) from small polysomes do not correlate with large polysomes. (B) Small polysome footprinting correlates highly with extensive experiment. (C) All 108 smORFs detected in large polysomes (blue) were also present in 198 from small polysomes (purple). (D) Three deep polysome profiling experiments reveal similar numbers of smORFs translated (Bar chart) and are highly coincident (Venn diagram). (E) 236 out of 274 smORFs show evidence of translation in at least one experiment, compared to 14151/17080 of standard ORFs. (F) Polysomal ribo-Seq detects 90% of smORFs with peptide evidence in S2 cells (52) or in all tissues (89) while rejecting less than 5%.

Small and large polysomes showed a marked difference in genome-wide ribosomal densities (Fig. 2A), compared to controls (Fig. S1C, D), suggesting that they contain different populations of mRNAs. Small polysomes contain mRNAs encoding long ORFs, but these have lower TE than when isolated from large polysomes (Table S2), confirming that they were bound by fewer ribosomes. As designed, our experiment also detected smORFs with translation signatures, and these were enriched in small polysomes (Table S3), which contained double and all the smORFs in large polysomes (Fig. 2C). The TE of smORFs from small polysomes is similar to the TE of long ORFs from large polysomes, indicating that smORFs are translated at similar levels to standard protein-coding ORFs (Fig. 1D).

Altogether 198 smORFs passed the cut-off values to be deemed translated in this initial deep polysome profiling experiment (Fig. 2C). This is nearly 72% of the smORFs transcribed in S2 cells in the total mRNA controls. To confirm and extend the catalogue of translated smORFs we repeated the experiment but exclusively sequenced small polysomes. This extensive small polysome profiling yielded double the number ORF-mapping reads obtained in the previous small polysome profiling (Table S4), and expanded the number of putatively translated smORFs to 234, which is 85% of smORFs we observe transcribed in S2 cells (Fig. 2D). The genome-wide distribution of ribosome densities in the two small polysome profiling experiments was strongly correlated (Fig. 2B), suggesting that deep polysome profiling is highly reproducible.

The majority of reads in both our experiments and previous ribosomal profiling consist of rRNA sequences liberated during footprinting (Table S4, (*13*)). Therefore we introduced rRNA-depletion beads during footprint extraction (see methods), which produced a marked improvement in the ratio of reads mapping to mRNAs and increased the total number of ORFs detected (Table S4, Fig. S2A) but did not expand our overall catalogue of putatively translated smORFs (Fig. 2D). The results of the three independent experiments are highly overlapping as 80% of putative translated smORFs were detected in all three datasets (Fig. 2D). By combining the three experiments we provide evidence that 236 smORFs are translated out of 274 transcribed in S2 cells (86%), which is very similar to the proportion of standard length protein-coding ORFs translated (82.8%) (Fig. 2E).

This high reproducibility might indicate that our screen for translated smORFs reached saturation, in other words, we have detected the near-total population of translated smORFs. To corroborate this, we compared our results with peptidomics data (Peptide Atlas, (*19*)). We detect translation of 96% of smORFs with peptidomic evidence in S2 cells (Fig. 2F), while only 3 smORFs whose translation we reject have similar evidence. This validates both our positive and negative results, confirming the thoroughness of polysomal ribo-Seq, which increases nearly 5-fold the number of smORFs with evidence of translation in S2 cells from 59 (proteomics) to 236. Our results seem extrapolable to *in vivo*, since we similarly detect as translated 90% of S2-transcribed smORFs with peptidomic evidence in any *Drosophila* tissue, while rejecting only 9 (10%) putative false negatives (Fig. 2F). Accordingly 202 smORFs detected as translated by us are transcribed in embryos, some 88 (40.2%) throughout the whole embryogenesis (Table S5). Overall, our 236 translated smORFs nearly doubles the previous number of smORFs with evidence of translation in any *Drosophila* tissue (147).

**FIGURE 3.**
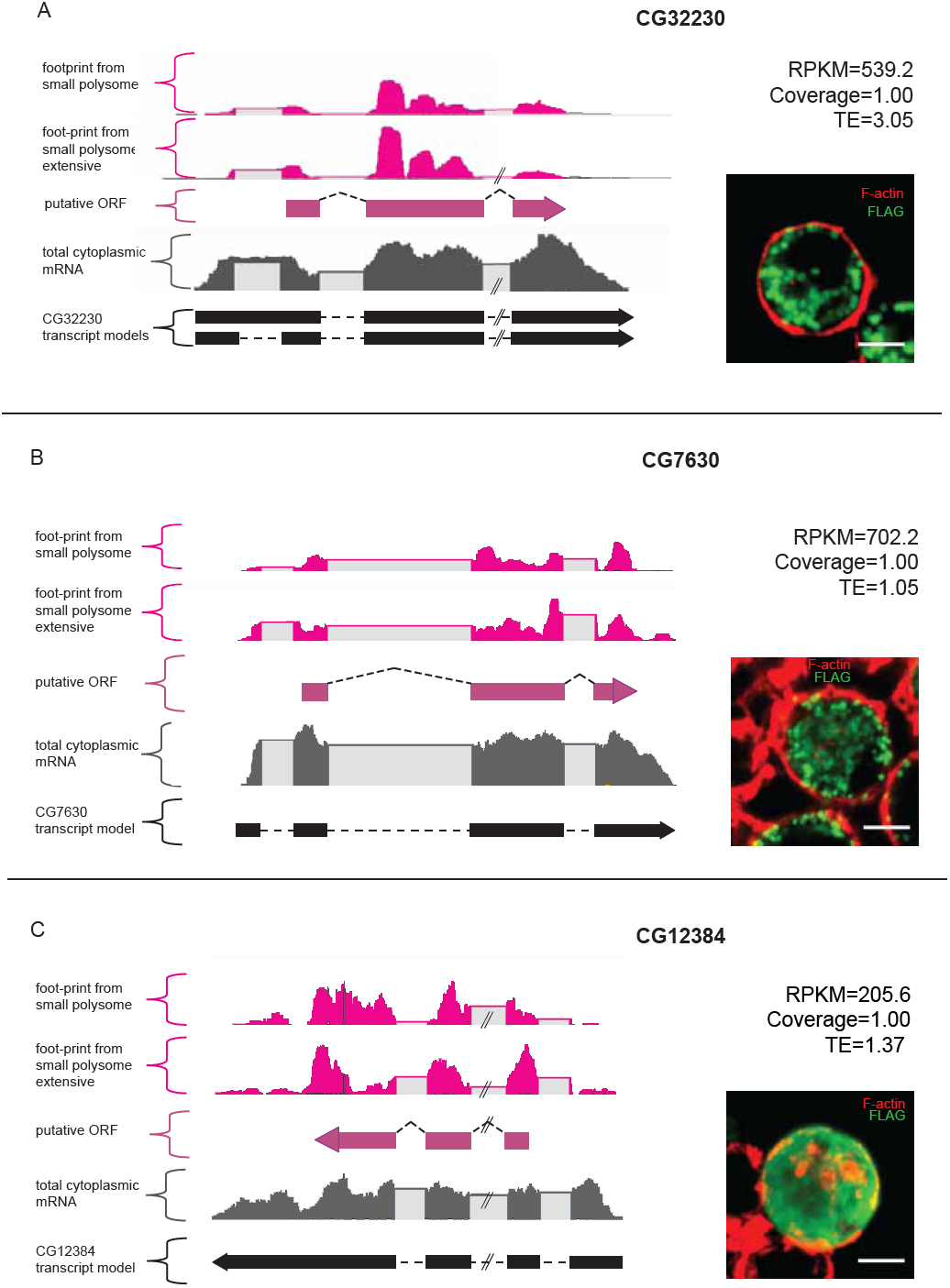
Validation of smORF translation by tagging assay. Ribosome footprints from small polysomes (pink) and mRNA reads (grey) mapped to smORFs, along with transcript and ORF models. Corresponding transfection assay in S2 cells are on right (green; FLAG antibody, red; F-actin stained with phalloidin, scale bars = 5 μm). (A) CG32230 (B) CG7630 (C) CG12384.

To validate the results of our smORF polysome profiling data we designed a peptide-tagging assay (Fig. S3A; methods). Transfection and staining of S2 cells with FLAG-tagged smORFs confirmed the translation of 18/19 smORFs with a range of translational indicators (Fig. 3A–C and S3; Table S6), indicating that even lower levels of translation can give rise to detectable smORF peptides.

**FIGURE 4.**
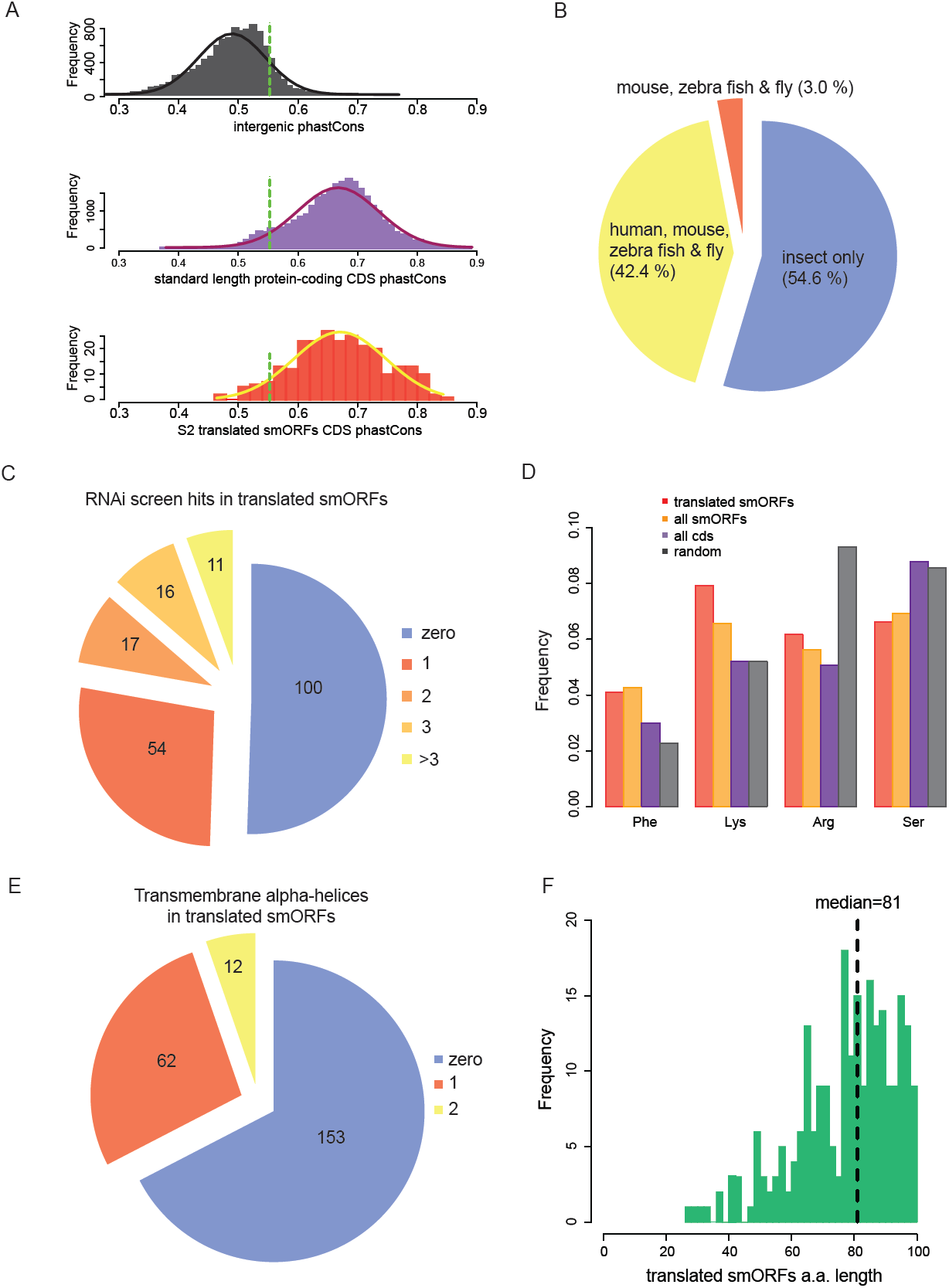
smORFs are conserved and functional. (A) Distribution of phastCons scores for intergenic regions, standard protein-coding CDSs and S2 cell-translated smORFs, with fitted normal curves. Green dotted lines indicate 90 percentile of intergenic phastCons score. (B) 45% of S2 cell-translated smORFs possess annotated vertebrate orthologs. (C) 50% of S2 cell translated smORFs show function in RNAi screens. (D) Relative abundance of particular amino acids in proteins, (expected, black; all ORFs, purple; all smORFs, yellow and translated smORFs, red). (E) Proportion of translated smORFs predicted to contain transmembrane α helices (F) Frequency distribution of peptide lengths of S2 cell-translated smORFs.

To assess the functionality of the translated smORF peptides, we studied their evolutionary sequence conservation as a proxy, since conservation usually implies selective retention of functional elements. We used phastCons (*20*), which measures conservation between twelve insect species. We observed the phastCons values in canonical long protein-coding sequences and in intergenic sequences and obtained a cut-off value of 0.55 separating them (10% FDR). 93.2% of S2-translated smORFs have phastCons score above this threshold (Fig. 4A), indicating a conservation level similar to that of canonical long ORFs, and hence, a similar level of functionality for the encoded peptides. We extended the time frame of comparative genomics up to mammals by compiling orthologous gene pairs between fly, zebra fish, mouse and human. Almost half of translated smORFs have annotated vertebrate orthologs (Fig. 4B). Such deep conservation is also indicative of function for the encoded peptides. In agreement, half of the translated smORF-encoding genes have revealed a function in S2 cell RNAi screens (*21*) (Fig. 4C).

We studied the amino acid usage in translated smORFs, compared to canonical long proteins and expected random usage (see methods and Fig. 4D). smORFs display a lower than random usage of arginine, which is a hallmark of translated proteins (*22*). However, smORFs display differential usage of several amino acids, characteristic of alpha-helices in canonical proteins (Fig. 4D). This finding was corroborated by an abundance of putative transmembrane alpha-helices, in about a third of translated smORFs (Fig. 4E) compared to the expected 20% (*23*). This is in agreement with similar findings in bacteria (*1*) and suggests that smORFs may represent a source of uncharacterised transmembrane peptides (Fig. S4C).

We note that the observed characteristics of translated smORFs (average size (Fig 4F, Fig. S4B), amino acid usage (Fig 4D, Fig. S4D) and abundance of putative transmembrane alpha-helices (Fig 4E, Fig. S4A, Fig. S4C) extends to the 555 annotated smORFs not transcribed in S2 cells. This might indicate that our results showing translation of 86% of transcribed smORFs could be extrapolated to this wider pool, possibly bringing the number of translated smORFs in *Drosophila* to ∼700. Since the estimated genome fraction that encodes smORFs in other organisms is similar to *Drosophila* (5%) (*5*), our results might suggest that hundreds of smORFs are translated in eukaryotic species.

In summary, deep polysome profiling increases 4-fold the number of smORFs in *Drosophila* S2 cells with evidence of translation from 59 to 236, and suggests that smORFs are translated in a similar proportion to canonical proteins longer than 100 aa. These smORFs have conservation levels similar to canonical proteins, and half have so far displayed functionality in genetic tests. Extrapolation of our results could indicate that hundreds of smORFs are translated in higher organisms and that their peptides might be active particularly in cell membranes. We surmise that a whole class of hundreds of small, membrane-associated peptides is awaiting characterisation and they could alter our understanding of many cellular and organismal processes.

## Acknowledgements

We thank Simon Morley and Ali Mumtaz for help with protocols, Claudio Alonso, Jose Ignacio Pueyo and Emile Magny for manuscript comments. This work was funded by a Wellcome Trust Fellowship (ref 087516).

## SUPPLEMENTARY MATERIALS

### MATERIALS AND METHODS

#### Tissue culture

S2 cells were grown under standard conditions in Schneiders media with 10% FBS.

#### Deep polysome profiling

S2 cells were treated with cycloheximide (Sigma) at 100 μg/ml for 3 min at RT before harvesting. Cells were pelleted for 8 min at 800 x g and washed with 1 X PBS with cycloheximide. Cells were resuspended in lysis buffer; 50 mM Tris-HCl pH8, 150 mM NaCl, 10 mM MgCl_2_, 1 mM DTT, 1% NP40, 100 µg/ml cycloheximide, Turbo DNase (Invitrogen), RNasin^®^ Plus RNase Inhibitor (Promega), cOmplete protease inhibitor (Roche). Nuclei were removed with a 5 min spin at 16,000 x g. Cycloplasmic lysates were loaded onto sucrose gradients and subject to ultracentrifugation. Gradients were pumped out, their absorbance at 254 nm plotted and fractionated. We used sucrose gradients to purify mRNAs in small polysomes, away from monosomes (80S), ribosomal subunits (40S, 60S) and large polysomes. Footprinting was performed overnight at 4°C with RNaseI (Invitrogen), stopped with SUPERase·In ^TM^ RNase inhibitor (Invitrogen) and then precipitated. mRNA from total cytoplasmic lysate, was purified using oligo (dT) Dynabeads (Invitrogen) and fragmented by alkaline hydrolysis. 28-34 nt ribosome footprints and 50-80 nt mRNA fragments were gel purified and prepared as previously described (*12*, *13*, *24*) for Next Generation Sequencing. Libraries were sequenced on Illumina HiSeq2000 and MiSeq machines with 50bp SingleEnd read protocol.

#### rRNA depletion

To generate ssDNA complementary to *Drosophila* rRNA, PCRs were performed to generate 500 and 1000 nt fragments of the rRNAs, using 5’ biotinylated reverse primers (sequences available on request). A 5’ biotinlyated oligo completementary to 2S rRNA, along with the PCR products were bound to magnetic streptavidin beads (Invitrogen). The second strand of the PCR fragments was washed away. Two rounds of 50 µl rRNA beads were used to deplete the all polysome sample prior to reverse transcription.

#### RT-PCR

RNA from sucrose gradient fractions was precipitated with isopropanol and 0.3 M NaCl. Resuspended pellets were treated with Turbo DNaseI (Ambion), extracted with phenol/chloroform and re-precipitated. 1µg RNA was used in RT reactions with MMLV reverse transcriptase (Promega). cDNA was subject to PCR with primers for specific mRNAs with Taq Polymerase (Qiagen).

#### Footprint sequence alignment

Sequences were aligned to FlyBase (Release 5.50) annotated transcript with previously developed bioinformatics analysis (*24*). Briefly, clipped and trimmed sequencing reads were aligned to an rRNA & tRNA reference using Bowtie short-read alignment program, discarding the rRNA and tRNA alignments and collecting unaligned reads. The unaligned reads were mapped to a genomic reference using the TopHat splicing-aware short-read alignment program. We only retained reads that were mapped to unique genomic locations. Alignments were accepted with up to two mismatches.

#### Footprint profile analysis

Profiles of ribosome footprints across a transcript were constructed by quantifying the number of footprint reads aligned at each position within the feature of interest. Two measurements were use to quantify translational level of a specific feature: ribosome density and coverage. Ribosome density was computed by scaling read counts for each feature-by-feature length and by the total number of genome-aligned reads (*12*). Footprint coverage estimated the percentage of each feature covered by ribosome footprints. We used the coverageBed command of the BEDTools program for computing coverage.

Translation efficiency (TE) was calculated as the ratio between the ribosome footprint density and the mRNA-seq read density in the window. As the TE score is not a reliable estimator at low expression levels, we computed a TE score only for those features that had significant mRNA expression above a randomized genomic background (p < 0.01). We placed no restrictions on the significance level for the ribosome coverage.

#### uORF identification

We identified canonical uORFs that initiate translation at AUG start codons based on the presence of an AUG followed by and in-frame stop codon within the annotated 5’-UTRs, using emboss getorf program. To exclude the possibility that the high ribosome occupancy observed in 5’-UTRs was due to the presence of upstream ORFs, we created a modified transcript that contained all regions except the putative uORFs for all our analysis on 5’-UTRs.

#### FlyBase smORFs analysis

We define smORFs genes as genes that encode for peptides of fewer that 100 amino acids long. Amongst FlyBase (release 5.50) protein-coding transcripts, 836 encode for peptides of less that 100 aa.

#### phastCons values

phastCons scores for 17,1317 alignment blocks were download from UCSC Genome Browser. We computed percentage overlap between the phastCons block and our feature of interest and estimated mean phastCons values for coding exon, intron, 5’- and 3’- UTR regions of all FlyBase protein-coding genes.

#### Peptide Atlas

List of peptide CDS coordinates with protein identifiers (FlyBase peptide ID) was downloaded from Peptide Atlas database (http://www.peptideatlas.org) and compared to FlyBase annotated smORF peptide sequences. 138 smORF proteins were hit by one or more sequenced peptide from the database.

#### Functional analysis of smORFs

Prediction of transmembrane alpha helices was performed using TMHMM (http://www.cbs.dtu.dk/services/TMHMM/). Vertebrate orthologs were identified by Ensembl (http://www.ensembl.org/index.html). In house perl scripts calculated amino acid composition of CDS and used all FlyBase transcipts as the random control. RNAi screen data was accessed through Flymine (http://www.flymine.org/) and GO term enrichment was calculated by Gorilla (http://cbl-gorilla.cs.technion.ac.il/) (*25*).

#### Cloning

The 5’-UTR and CDS of putative smORFs were cloned by PCR from S2 cell cDNA into pENTR^TM^ /D-TOPO^®^ (Invitrogen) and then into pAWF (http://emb.carnegiescience.edu/labs/murphy/Gateway%20vectors.html#), whose ATG start codon was mutated to GCG by site-directed mutagenesis.

#### Transfections and microscopy

250,000 S2 cells were plated on acid-treated coverslips and transfected with 1 ug plasmid DNA using Xtreme Gene HP (Roche). After 48 h, cells were fixed for 20 min with 4% formaldehyde and washed with 1 X PBS, 0.1 % Triton X-100 (PBS-T). Transfections with FLAG-tagged constructs were blocked with PBS-T 2% w/v BSA before immunostaining with primary mouse anti-FLAG M2 antibody (SIGMA) at 1/1000 and secondary anti-mouse FITC (Jackson) at 1/400. All transfected cells were incubated for 30 min with Rhodamine-Phalloidin (Invitrogen) to highlight F-actin then 10 min with Hoechst (SIGMA) for nuclei staining and mounted with Vectashield (Vector Labs). Imaging was conducted using a Zeis 63X oil Plan Apochromat on the LSM510 metAxioskop2.

#### SUPP. TABLES

**SUPPLEMENTAL TABLE S1.** Summary across experiments of 90 percentile for 3’-UTR ribosomal density and coverage.

| 90 percentile | Small polysomes | Large polysomes | Extensive small polysomes | -rRNA all polysomes | Mean |
| --- | --- | --- | --- | --- | --- |
| RPKM 3'-UTR | 5.54 | 7.67 | 8.95 | 8.64 | 7.70 |
| Coverage 3'-UTR | 0.499 | 0.471 | 0.718 | 0.745 | 0.57 |

**SUPPLEMENTAL TABLE S2.**
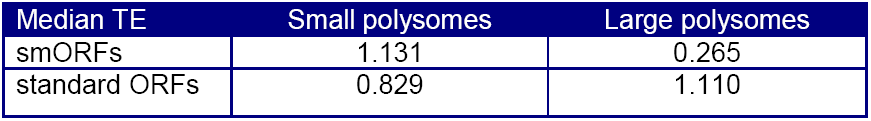
Median translational efficiency for smORFs and standard ORFs in small and large polysomal fractions.

| Median TE | Small polysomes | Large polysomes |
| --- | --- | --- |
| smORFs | 1.131 | 0.265 |
| standard ORFs | 0.829 | 1.110 |

**SUPPLEMENTAL TABLE S3.** Specific enrichment of smORF translation signal in small polysomes compared to large polysomes. Chi squared test on the number of ORFs translated in small and large polysomal fractions (p=0.00039).

|  | Translated standard protein-coding ORFs | Translated smORFs |
| --- | --- | --- |
| Small polysomes | 10382 | 198 |
| Large polysomes | 8717 | 108 |
| Ratio | 1.19 | 1.83 |

**SUPPLEMENTAL TABLE S4.** Summary of sequencing experiments. Number of reads from each experiment, number that are left after removal of rRNA and tRNA contaminants, as well as proportion of reads that are unique matches and number of reads mapping to CDS regions of the genome.

| Experiment | Raw reads | Reads that pass clipping and trimming | After removal of rRNA and tRNA | Tophat mapped reads | Unique match reads | Reads that map to ORFs |
| --- | --- | --- | --- | --- | --- | --- |
| Small polysomal footprint | 99,973,197 | 93,838,021 | 6,250,297 (6.66%) | 5,498,415 (87.97%) | 3,748,512 (68.17%) | 2,870,975 (76.59%) |
| Large polysomal footprint | 67,230,885 | 59,191,477 | 3,579,490 (6.05%) | 2,625,211 (73.34%) | 2,371,751 (90.34%) | 1,906,050 (80.36%) |
| Total cytoplasmic mRNA 1 | 3,737,677 | 3,296,829 | 2,249,793 (68.24%) | 2,006,335 (89.17%) |  | 1,619,013 (80.7%) |
| Total cytoplasmic mRNA 2 | 8,698,975 | 4,374,205 | 2,911,316 (66.56%) | 2,559,302 (87.91%) |  | 1,956,992 (76.47%) |
| Small polysomal footprint extensive | 189,631,476 | 188,066,263 | 20,008,723 (10.64%) | 9,160,610 (45.78%) | 8,133,854 (88.79%) | 5,904,132 (72.59%) |
| -rRNA all polysomes footprint | 14,092,100 | 12,759,694 | 4,144,111 (32.48%) | 3,190,896 (77.0%) | 2,972,003 (93.14%) | 2,355,257 (79.25%) |

**SUPPLEMENTAL TABLE S5.** Summary of tagged smORFs. Details of the deep polysome profiling results and transfection translation assay results for the FLAG-tagged smORFs, with RPKM and coverage values from the small polysome experiment.

| Number of Embryo Stages | Number of smORFs (%) |
| --- | --- |
| 12 | 88 (40.2%) |
| 11 | 17 (7.8%) |
| 10 | 11 (5.0%) |
| 9 | 18 (8.2%) |
| 8 | 10 (4.6%) |
| 7 | 9 (4.1%) |
| 1-6 | 49 (22.4%) |
| 0 | 17 (0.078%) |
| **TOTAL** | **219** |
| NA | 17 |

**SUPPLEMENTAL TABLE S6.** Number of translated smORFs expressed throughout embryonic stages of *Drosophila melangogaster*.

| smORF | FBpp | RPKM | coverage | TE | aa length | FLAG distribution | ORF phasCon |
| --- | --- | --- | --- | --- | --- | --- | --- |
| CG32230 | FBpp0070039 | 539.2 | 1.00 | 3.05 | 83 | Punctate | 0.54 |
| CG7630 | FBpp0074979 | 702.2 | 1.00 | 1.05 | 90 | Punctate | 0.64 |
| CG12384 | FBpp0087252 | 205.6 | 1.00 | 1.37 | 96 | Dispersed cytoplasmic | 0.71 |
| CG33170 | FBpp0289155 | 84.2 | 1.00 | 0.75 | 71 | Mixed | 0.60 |
| CG14482 | FBpp0086062 | 600.0 | 1.00 | 1.09 | 57 | Punctate | 0.72 |
| CG33774 | FBpp0091036 | 115.3 | 1.00 | 1.13 | 40 | Dispersed cytoplasmic | 0.73 |
| CG44242 | FBpp0308229 | 152.9 | 0.97 | 1.75 | 70 | Punctate | 0.66 |
| CG32267 | FBpp0073001 | 82.5 | 0.97 | 1.13 | 49 | No signal | 0.70 |
| CG15456 | FBpp0077015 | 10.1 | 0.65 | 0.95 | 95 | Dispersed cytoplasmic | 0.68 |
| CG33199 | FBpp0087769 | 95.5 | 1.00 | 1.17 | 79 | Punctate | 0.59 |

**SUPPLEMENTAL FIGURE 1.**
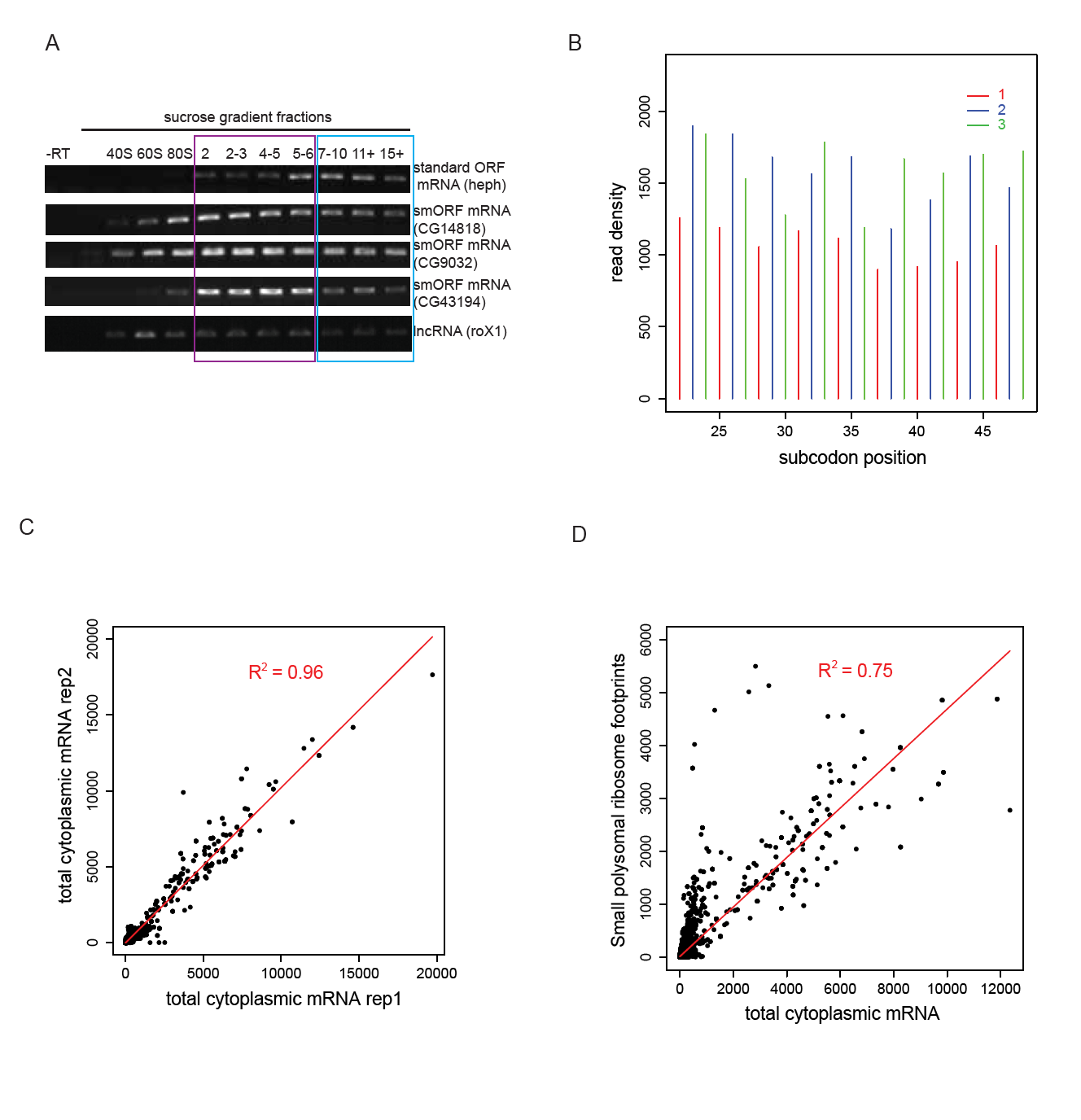
(A) RT-PCR of RNA recovered from sucrose gradient fractions for one standard ORF mRNA (heph), three smORF mRNAs (CG14818, CG9032 and CG43194) and roX1 lncRNA, with -RT control. (B) Read density plot showing phasing of ribosome footprinting reads in triplets corresponding to codons (small polysomes). (C) Read densities (RPKM) from two biological replicates of control total cytoplasmic mRNA exhibit very high correlation (R^2^=0.96). (D) Ribosome footprinting densities from small polysomes are substantially different from read densities from control total cytoplasmic mRNA (R^2^= 0.75).

**SUPPLEMENTAL FIGURE 2.**
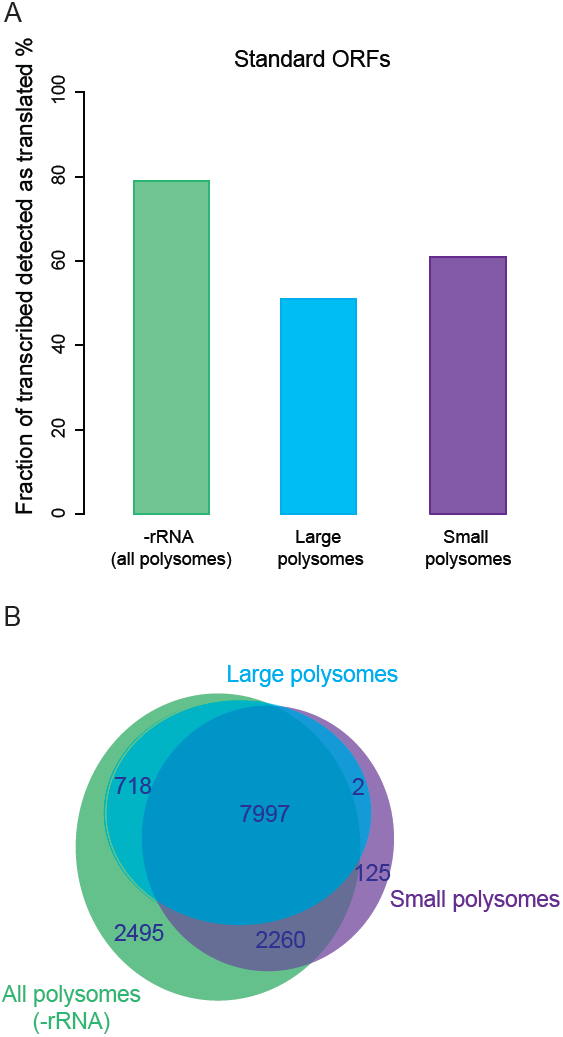
Deep polysome profiling experiments of all (-rRNA), large and small polysomes reveal proportion of standard protein-coding ORFs translated (A) and overlap between experiments (B).

**SUPPLEMENTAL FIGURE 3.**
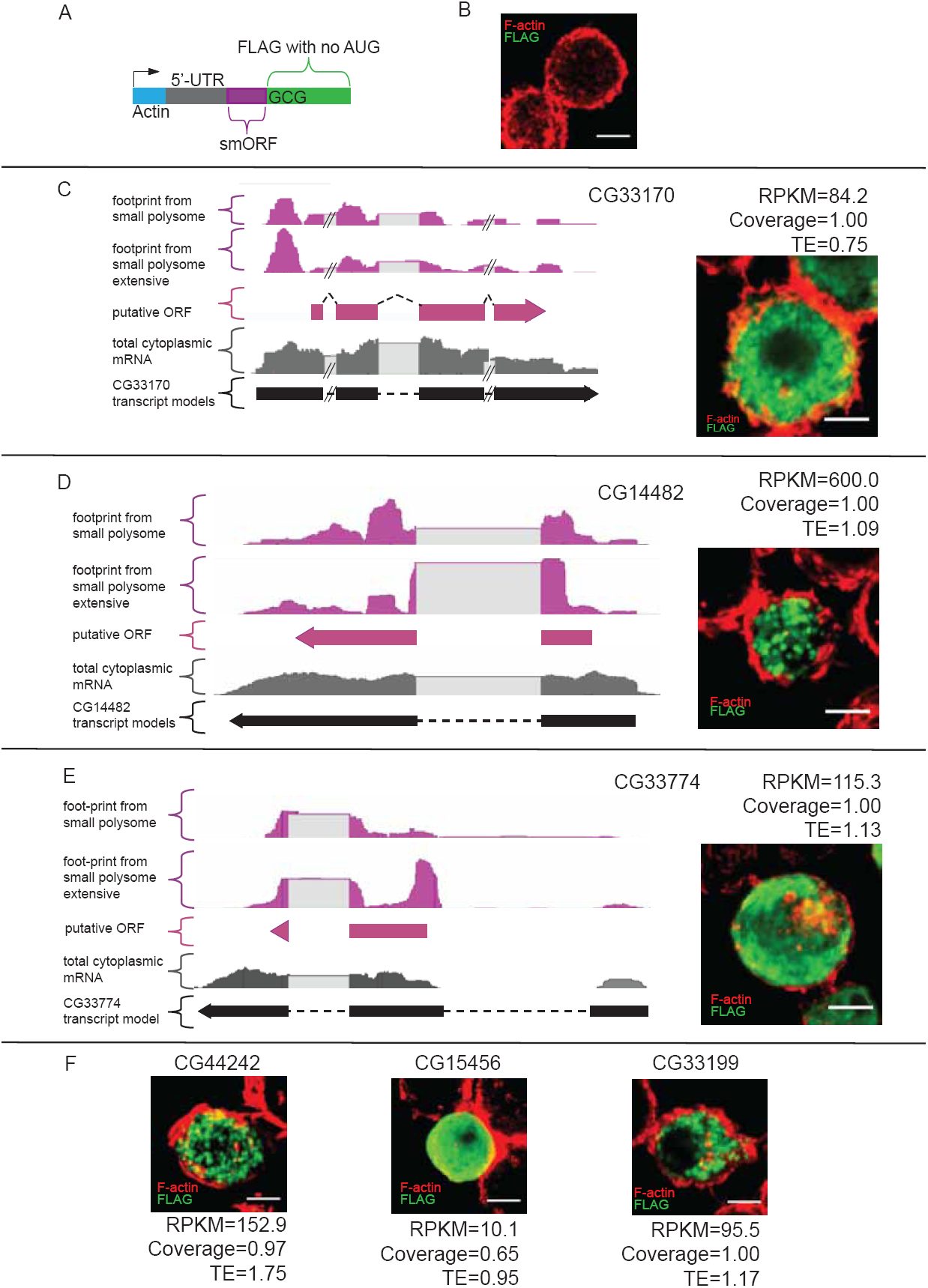
(A) Schematic of transfection construct into which smORF 5’-UTRs and ORFs (no stop codon) were cloned under Actin promoter, fused to C-terminal FLAG tag, with AUG start codon mutated to GCG, in frame with smORF. (B) Transfection negative control, plasmid with no ORF (nor AUG) cloned in. Ribosome footprints from small polysomes and total cytoplasmic mRNA reads mapped to smORFs, with transcript and ORF models (C) CG33170, (D) CG14482 and (E) CG33774. Transfection translation assay in S2 cells for smORFs; (C) CG33170, (D) CG14482, (E) CG33774, (F) CG44242, CG15456, CG33199, (green; FLAG antibody, red; F-actin stained with phalloidin, scale bars = 5 μm).

**SUPPLEMENTAL FIGURE 4.**
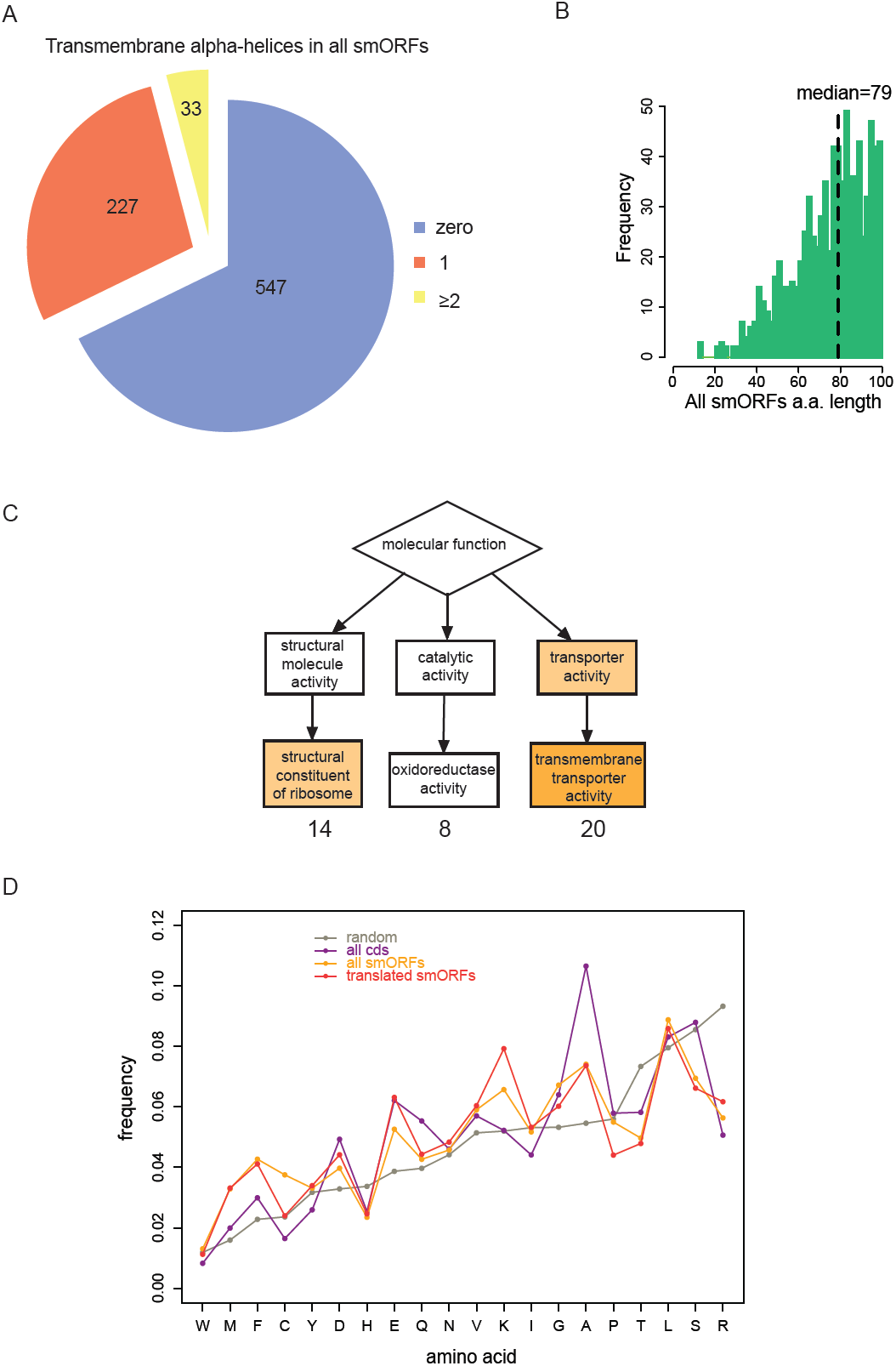
(A) Proportion of all 829 annotated smORFs predicted to contain transmembrane α helices (according to TMHMM). (B) Frequency distribution of peptide lengths of all smORFs. (C) Enrichment of GO terms (GOrilla) within translated smORFs compared to translated standard protein-coding ORFs in S2 cells; structural consitituents of ribosome (p=1.25E-4), oxidoreductase activity, transmembrane transporter activity (p=7.35E-6). (D) Relative abundance of all amino acids in proteins, (random, blue; all ORFs, green; all smORFs, purple, and translated smORFs, red).

## References

1. M. R. Hemm, B. J. Paul, T. D. Schneider, G. Storz, K. E. Rudd, Mol Microbiol 70, 1487 (Dec, 2008).

2. M. A. Basrai, P. Hieter, J. D. Boeke, Genome Research 7, 768 (August 1, 1997, 1997).

3. J. P. Kastenmayer et al., Genome Res. 16, 365 (March 1, 2006, 2006).

4. K. Hanada et al., Proc Natl Acad Sci U S A 110, 2395 (Feb 5, 2012).

5. E. Ladoukakis, V. Pereira, E. G. Magny, A. Eyre-Walker, J. P. Couso, Genome Biol 12, R118 (2011).

6. M. C. Frith et al., Plos Genetics 2, 515 (Apr, 2006).

7. S. A. Slavoff et al., Nat Chem Biol 9, 59 (Jan, 2013).

8. W. F. Burkholder, I. Kurtser, A. D. Grossman, Cell 104, 269 (Jan 26, 2001).

9. M. I. Galindo, J. I. Pueyo, S. Fouix, S. A. Bishop, J. P. Couso, Plos Biology 5, 1052 (May, 2007).

10. E. G. Magny et al., Science 341, 1116 (Sep 6, 2013).

11. L. X. Shi, W. P. Schroder, Biochim Biophys Acta 1608, 75 (Feb 15, 2004).

12. N. T. Ingolia, S. Ghaemmaghami, J. R. Newman, J. S. Weissman, Science 324, 218 (Apr 10, 2009).

13. N. T. Ingolia, L. F. Lareau, J. S. Weissman, Cell 147, 789 (Nov 11, 2011).

14. G. L. Chew et al., Development 140, 2828 (Jul, 2013).

15. M. Guttman, P. Russell, N. T. Ingolia, J. S. Weissman, E. S. Lander, Cell 154, 240 (Jul 3, 2013).

16. B. A. Wilson, J. Masel, Genome Biol Evol 3, 1245 (2011).

17. I. Schneider, J Embryol Exp Morphol 27, 353 (Apr, 1972).

18. Y. Arava et al., Proc Natl Acad Sci U S A 100, 3889 (Apr 1, 2003).

19. F. Desiere et al., Genome Biol 6, R9 (2005).

20. A. Siepel et al., Genome Res 15, 1034 (Aug, 2005).

21. E. E. Schmidt et al., Nucleic Acids Res 41, D1021 (Jan, 2012).

22. J. L. King, T. H. Jukes, Science 164, 788 (May 16, 1969).

23. A. Krogh, B. Larsson, G. von Heijne, E. L. Sonnhammer, J Mol Biol 305, 567 (Jan 19, 2001).

24. N. T. Ingolia, G. A. Brar, S. Rouskin, A. M. McGeachy, J. S. Weissman, Nat Protoc 7, 1534 (Aug, 2012).

25. E. Eden, R. Navon, I. Steinfeld, D. Lipson, Z. Yakhini, BMC Bioinformatics 10, 48 (2009).

